# Ion-Concentration Gradients During Synaptic Input Increase the Voltage Depolarization in Dendritic Spines

**DOI:** 10.1101/2023.08.03.551788

**Authors:** Florian Eberhardt

## Abstract

The cable equation is key for understanding the electrical potential along dendrites or axons, but its application to dendritic spines remains limited. Their volume is extremely small so that moderate ionic currents suffice to alter ionic concentrations. The resulting chemical-potential gradients between dendrite and spine head lead to measurable electrical currents. The cable equation, however, considers electrical currents only as result of gradients in the electrical potential. The Poisson-Nernst-Planck (PNP) equations allow a more accurate description, as they include both types of currents. Previous PNP simulations predict a considerable change of ionic concentrations in spines during an excitatory postsynaptic potential (EPSP). However, solving PNP-equations is computationally expensive, limiting their applicability for complex structures.

Here, we present a system of equations that generalizes the cable equation and considers both, electrical potentials and time-dependent concentrations of ion species with individual diffusion constants. Still, basic numerical algorithms can be employed to solve such systems. Based on simulations, we confirm that ion concentrations in dendritic spines are changing significantly during current injections that are comparable to synaptic events. Electrical currents reflecting ion diffusion through the spine neck increase voltage depolarizations in the spine head. Based on this effect, we identify a mechanism that affects the influx of Ca2+ in sequences of pre- and postsynaptic action potentials. Taken together, the diffusion of individual ion species need to be taken into account to accurately model electrical currents in dendritic spines. In the future the presented equations can be used to accurately integrate dendritic spines into multicompartment models to study synatptic integration.

## 1 Introduction

Synapses on dendritic spines are the major recipients of excitatory input to pyramidal neurons [21]. During activation of the synapse, synaptic currents depolarize the spine head [1]. The synaptic currents are then transmitted to the parent dendrite, and the collective input to the neuron gets integrated along the dendritic tree [6]. However, electric signals can also be transmitted in the opposite direction. A back-propagating action potential (bAP) is generated at the axon hillock and travels up the dendritic tree and finally depolarizes the cell membrane in dendritic spines [28]. Additionaly, dendrititc spikes, namely dendritic sodium, calcium, or N-methyl-D-aspartate (NMDA) spikes exist. In contrast to somatic action potentials, these local regenerative potentials can be triggered in the dendritic tree [18]. While spines can compartmentalize electrical and chemical signals evoked by synaptic input [3, 10, 15], postsynaptic signals (bAPs or dendritic spikes) are assumed to invade the spines without attenuation [19, 31]. The integration of signals related to pre- and postsynaptic activity is of great interest, as the activation of voltage-dependent NMDA receptors in dendritic spines can induce synaptic plasticity through calcium influx [2, 22]. To study electric signals, propagating along a dendrite the cable equation can be used [26]. The assumptions of cable theory are well-justified for dendrites, but unfortunately the formalism fails when applied to very small structures such as dendritic spines [24].

The cable equation models the electric potential along a dendrite or an axon by its membrane capacitance, intracelluar resistance and membrane resistivity. However, the ionic composition of the intracellular electrolyte is assumed to be constant and currents induced by the diffusion of ions are ignored. Because of the small spine volume, e.g. on average 0.02*μm*^3^ on pyramidal cells [12], the ion-concentrations can easily change in spines [17]. A typical pyramidal cell spine head contains 1.2 ·10^5^ sodium ions at a concentration of 10 mmol. During an EPSP with a synaptic current of 23 pA [10] that lasts for 12 ms [1], 14.0 ·10^5^ ions enter the cell. The number of sodium ions entering the spine head during a synaptic event is therefor significantly higher, than the number of sodium ions located inside the spine head at rest. The resulting concentration change in the spine head persists for several milliseconds, even in the case of small ions. The decay time for sodium is approximately 8*ms* [30] for a typical pyramidal cell spine with a neck diameter of *r*_*Neck*_ = 50*nm*, a neck lenght of *l*_*Neck*_ = 500*nm*, a head readius of *r*_*Head*_ = 250*nm* and a diffusion constant of *D*_*Na*_ = 0.5*μm*^2^*/ms*, [17, 27]. The diffusion of the ions will induce electric currents across the neck that are not captured by cable theory [24]. Therefor, Poisson-Nerns-Planck (PNP) equations [20] have been used to accurately model the electric function of dendritic spines [5]. But especially in time dependent systems a numerical implementation of the PNP formalism is non-trivial and computationally expensive. Consequently only small systems with simple geometry are usually studied [7, 23].

Here we present an extension to cable theory derived from the Nernst-Planck-equations [9] for dendritic spines. This system of equations can be considered as a generalized form of the cable equation. However, it considers not only the electric potential, but also the time dependent concentrations of various ion species with individual diffusion constants. Still, basic numerical algorithms based on finite differences are sufficient to find solutions even in systems that include multiple ion-species.

Based on computer simulations, we find that a depolarization of the parent dendrite will invade spine heads without attenuation, in agreement with previous experimental results [10]. We also confirm that synaptic currents alter ion concentrations in the spine head. For example, the sodium concentration increases by 280% for spine parameters found in [10]. Interestingly, the spine head depolarization is larger than expected from simple electrical models of spines. This shows that the depolarization cannot be explained by the synaptic current and the electrical resistance of the spine neck alone, but ion diffusion needs to be taken into account, too. In sequences, where postsynaptic activity follows presynaptic activity, this effect increases the influx of calcium into dendritic spines through voltage dependent ion-channels. The system of equations presented here includes all parameters relevant in cable theory. Thereby, the equations can be used in the future to extend multi-compartment models, and to accurately study effects of dendritic spines in synaptic integration.

## 2 Methods

The cable equation [25] describes how the temporal evolution of the electrical potential Φ along a passive neuronal process with length *l* and varying radius *a* depends on the membrane currents, ohmic axial currents, and on the membrane capacitance.

In this section we extend this fundamental equation to include currents induced by the diffusion of intracellular ions. Conceptuatlly similar derivavtion can be found for example in [14] to derive fractional cable equations for dendrites. The derivation is based an analogy between the electrical potential Φ and the chemical potential *μ* and on the Nernst-Planck equation [9]. The Nernst-Planck equation reads as

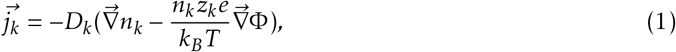

where *j*_*k*_ describes the particle current density of an ion-species *k* with density *n*_*k*_ as a result of diffusion and an electric field. *D*_*k*_ denotes the diffusion constant, *z*_*k*_ the charge number, *e* the elementary charge, *k*_*B*_ the Boltzmann constant and *T* the temperature.

In the following we consider a 1D-case with radial symmetry and without radial dependence of the electrical field and the chemical potential gradient. To transform an particle currents into electrical currents one can simply multiply the particle currents *j*_*k*_ by *z*_*k*_*e*. The electrical axial current of an ion-species *k* along a cylinder with cross-section *A* = *πa*^2^ is therefor

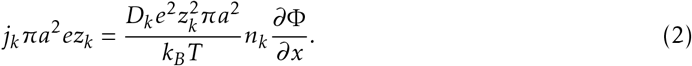

We now define

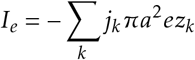

as the total axial electric current and

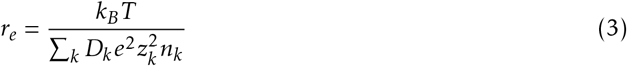

as the electrical resistivity. Together wis eq. (2) this leads to

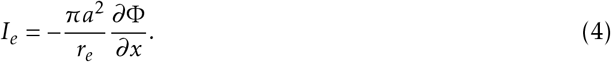

The chemical potential *μ*_*k*_ of an ion-species *k* if given by

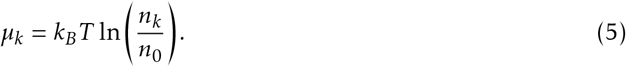

The gradient of the chemical potential is simply

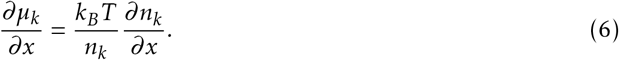

According to the Nernst-Planck equation the axial electrical current induced by the diffusion of ion species *k* can be computed as

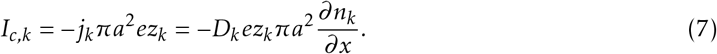

Next we define the diffusional resistivity

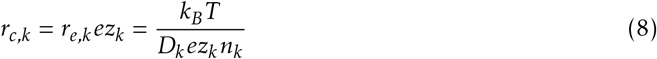

in analogy to the electrical resistance *r*_*e*_ and insert it into eq. (7). Together with eq. (6) this leads to

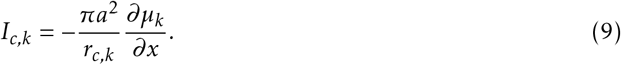

To derive the cable equation one balances the capacitive current and all currents that enter (or leave) a cable segment of finite length Δ*x. c*_*m*_ denotes the specific membrane capacitance. For simplicity we assume a system without any membrane currents.

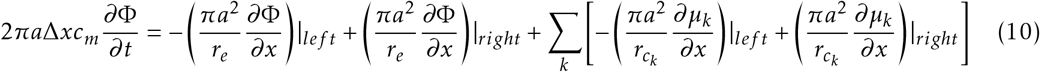

As described in [11] we divide this equation by 2*πa*Δ*x* and take the limit Δ*x →*0. We further replace the gradient of the chemical potential using eq. (6). Compared to its standard form this leads to a cable equation with an additional diffusion term.

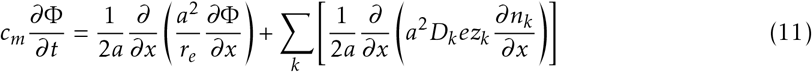

The same derivation can be carried out for the chemical potential to describe temporal changes in the ionic concentrations along the cable. First we sum over all particle currents entering or leaving the volume:

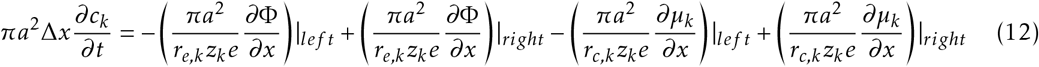

Then we divide the above equation by *πa*^2^Δ*x* and take the limit Δ*x* → 0 again, to arrive at

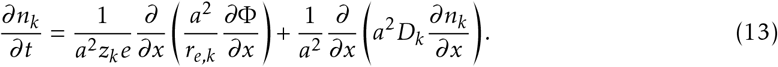

Equations (11) and (13) together with adequate boundary conditions, i.e. initial conditions uniquely determine the solution. By setting 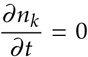 one recovers the well known standard form of the cable equation.

Numerically, equation (11) and (13) can be simply solved by an explicit forward timestepping algorithm based on finite differences. Stability of the solution can be achieved at low spatial and high temporal resolution. To avoid the independent drifting of the electrical potential and concentration variables eq. (11) should be replaced by a capacitor Φ = *c*_*m*_*πa*^2^Δ*xΣ_k_ z*_*k*_*en*_*k*_ + Φ_0_ and an additional fixed background charge to ensure electro-neutrality. Both approaches are equivalent, because

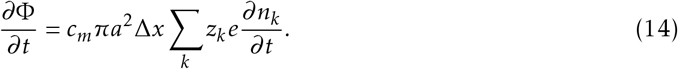

The spatial grid of the computational domain can be seen in Fig. 1A. The region with larger radius *a* on the right side represents the dendritic end where we apply the Dirichlet boundary conditions 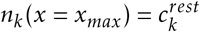 and Φ(*x* = *x*_*max*_) = Φ^*rest*^. On the synaptic end on the left we apply the Neumann boundary conditions, *∂*Φ*/∂x* = 0, *∂n*_*K*_ */∂x* = 0, *∂n*_*Cl*_*/∂x* = 0 and *∂n*_*Na*_*/∂x* = −*I*_*in*_*/*(*D*_*Na*_*ez*_*Na*_*πa*^*2*^) to model the sodium influx. Here, *I*_*in*_ denotes the input current on the synaptic end of the computational domain (red line in Fig. 1A).

**Figure 1:**
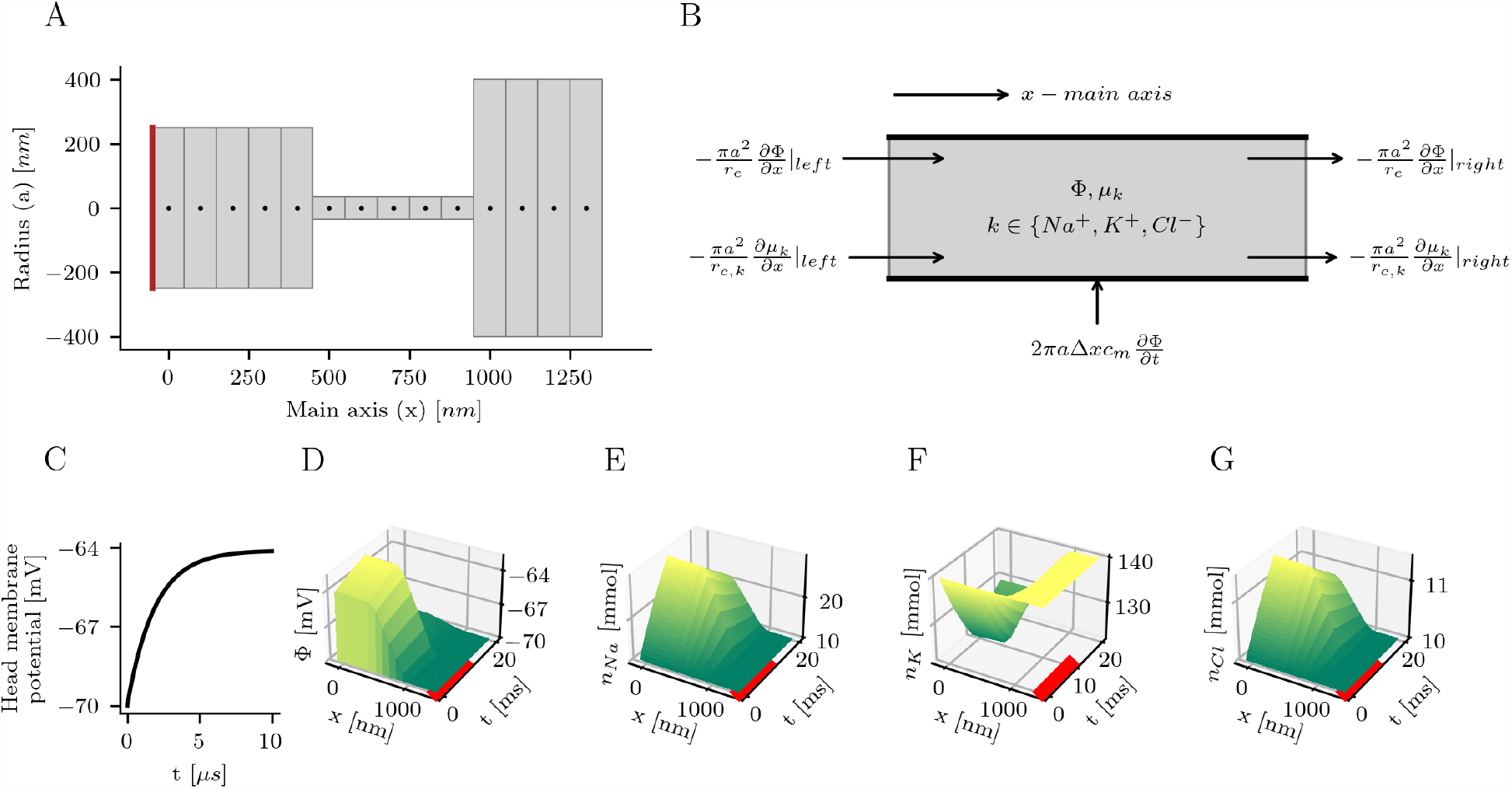
Simulation of an idealized dendritic spine based on parameters found in [10] A) The solution is computed on a lattice with a finite-difference algorithm. The one-dimensional spatial lattice is indicated by the black dots. Neumann boundary conditions are used to inject a sodium current to the spine head on the left side (red line) and to establish reflecting boundaries for potassium and chloride, and the elctrical potential. On the right, i.e., dendritic end ion concentrations are set to constant values to ensure a smooth transition between spine and dendrite (Dirichlet boundary conditions). B) In cable theory the axial electric currents are computed from the gradient of the electric potential along the cable. By an analogy between the electric potential and the chemical potential the same formalism can be developed for the diffusion of ions. C-E) We simulate the membrane potential and the concentrations of sodium, potassium and chloride on the spatial lattice with forward time stepping. A 25 *pA* current of sodium ions gets injected for 10 *ms* (red bar). C) The time course of the membrane potential shows a rapid charging of the membrane capacitor within a few microseconds. D) The injected current depolarizes the spine head by several millivolts. The voltage drops almost completely across the spine neck. After the fast charging of the capacitor, the membrane voltage continues to slowly further increase. During the current injection, sodium E) and chloride F) concentrations are increasing while the potassium G) concentrations is decreasing in the simulated spine.

## 3 Results

Because of the small size of dendritic spines, even moderate ion influxes are likely to have a strong influence on the spatio-temporal dynamics of the ionic concentrations (*n*_*k*_) within spines. Through the diffusion of ions, the concentration gradients will induce additional electrical currents. A system of coupled partial differential equations allows one to calculate the temporal evolution of the concentration of the three major ion species, i.e., sodium, potassium and chloride, along a cable.

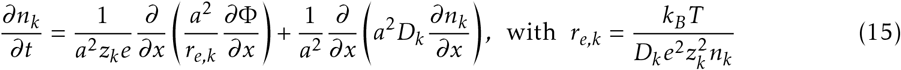

The electric potential can be computed from the summed charge and the specific membrane capacitance *c*_*m*_.

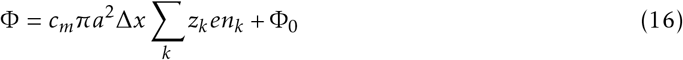

For a detailed derivation of these equations see the methods section. Their solution is obtained using a finite-difference grid, whose spatial x-axis can be seen in Fig. 1A, together with the radius parameter *a* of eq. (15). The wider region on the left indicates the spine head. We also include a section of the parent dendrite on the right. Spine head and dendrite are connected by a narrow spine neck. Neumann boundary conditions are used to inject a sodium current to the spine head on the left side and to establish reflecting boundaries for potassium and chloride, and the elctrical potential. On the right, i.e., dendritic end ion concentrations are set to constant values to ensure a smooth transition between spine and dendrite (Dirichlet boundary conditions). The initial values for Φ and *n*_*k*_ are set at the beginning of the simulation. The diffusion constants are 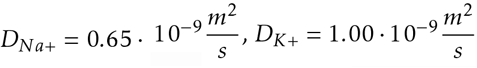 and 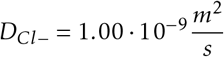. The values are chosen in agreement with [27]

### Ion concentrations in the spine head change during current injection

Recently, voltage compartmentalization in dendritic spines was measured for the first time in vivo [10]. Spines on pyramidal neurons in somatosensory cortex were found to compartmentalize the membrane voltage at more than 5*mV* on average. At an average photostimulation currrent of 23 pA the neck resistance was estimated to be approximately 230 *M*Ω. We choose the first simulation parameters accordingly, to reproduce these results. The spine neck diameter is set to 70 *nm* and a neck length to 500 *nm*. The neck resistance is estimated from the ion concentrations at rest to be 230 *M*Ω, using eq. 3. The head volume is set to 0.01 *μm*^3^ to match the typical volume of the intracellular space of spines from hippocampal pyramidal cells, excluding organelles and cytoskeleton [12]. Then we inject a constant influx of sodium ions that leads to an electric current of 25*pA* for a time of 10*ms*. The duration of the current injection is choosen in agreement with the average time of synaptic opening of 12*ms* in basal dendrites of L5 pyramidal neurons in acute brain slices [1].

The simulations show that the charging of the membrane capacitance is fast and only takes several microseconds. This agrees with [17]. The near instantaneous charging of the membrane capacitance is also consistent with the extremely small membrane capacitance of the spine head. After the rapid charging, the membrane of the head membrane is depolarized by aroung 6 *mV*. The head’s depolarization increases to 7.2 *mV* during the full 10 *ms* of current injection. The electrical potential is almost constant along the spatial axis within the head and drops almost linearly along the neck. The ion concentrations are significantly changing during current input. After 10 *ms* the sodium concentration increases from 10.0 *mmol* to 28.6 *mmol*, chloride from 10.0 *mmol* to 11.4 *mmol* and the potassium concentration drops from 140.0 *mmol* to 122.0 *mmol*. When the current injection ends, the electric potential drops within few microseconds to a value of −68.8 *mV*, close to the resting voltage of 70 *mV*, as the membrane capacitance discharges. Then the electric potential and the concentrations decay slowly towards their initial values. The decay time constant of sodium for the chosen simulation parameters can be estimated to 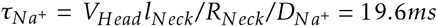 theoretically [30]. *R*_*Neck*_ is the total resistance of the spine neck. In the simulation we find a decay time constant of 19.2 *ms* consistent with the predicted value.

Experimental measurements usually show a high variance of the EPSP peak voltages in spines [10], and other studies also measure higher voltages or currents [1, 16] as assumed for the simulation of Fig. 1. This is in agreement with spines existing in various shape and size. The spine morphology gets usually quantified by the head volume, the neck width and the neck length. We simulate the potential and the ion concentrations for different shape parameters and different strength of input current (Fig. 2). As expected the neck diameter is the main determinant of the neck resistance (if the length is identical). The depolarization of the spine head is stronger in spines with thin necks. Moreover, we find that in smaller spine heads the concentrations changes are stronger, than in large spines (compare spine B with spine E and spine C with spine D). Especially in spines with small head volumes and thin necks (Fig. 2, spine B) the the ion concentrations change considerably. The sodium concentration increased from 10 mmol to more than 70 mmol, while the potassium concentration dropped accordingly.

**Figure 2:**
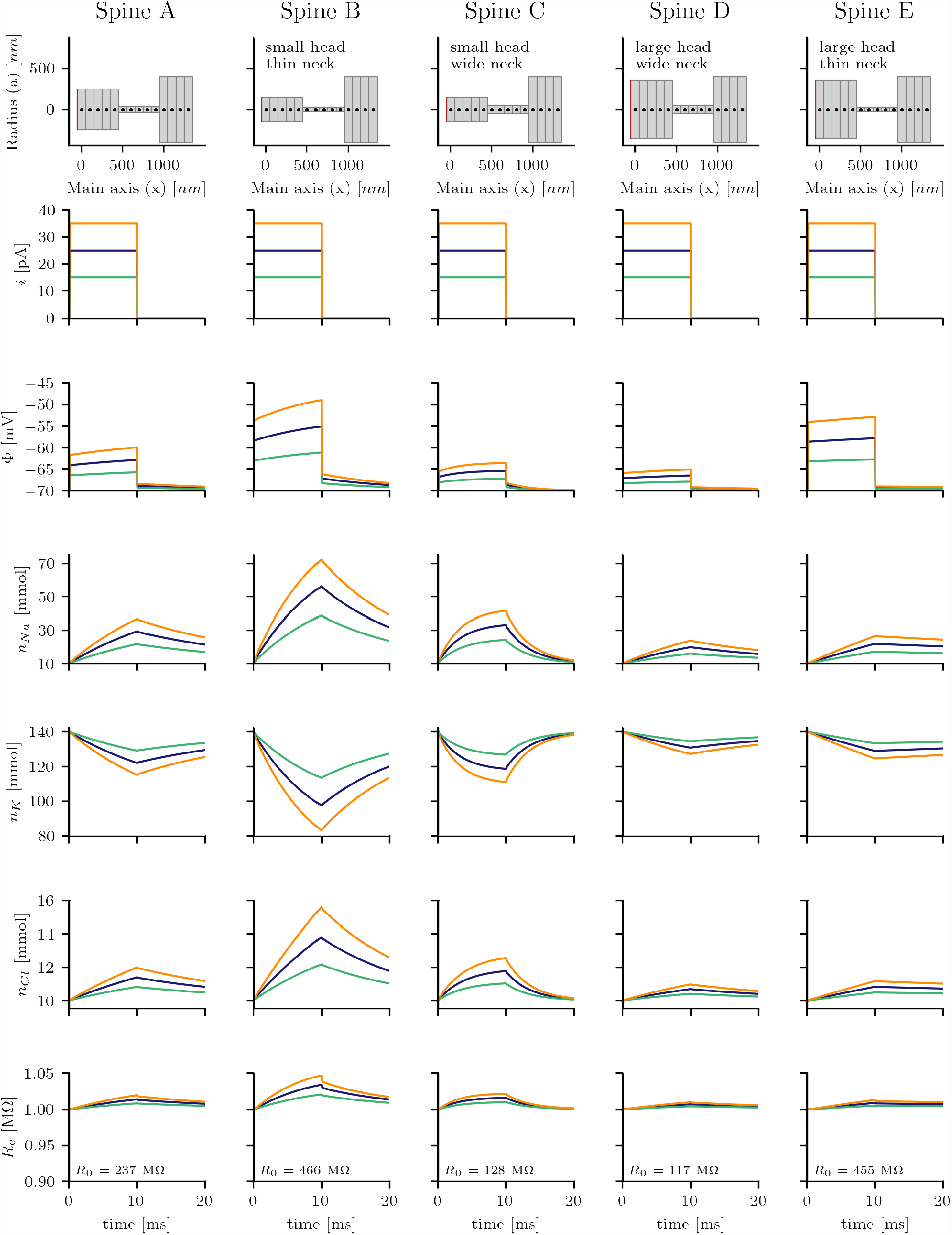
Changes in the membrane voltage and the ion concentrations of sodium, potassium and chloride related to the current injections (step current from *t* = 0 to *t* = 10*ms*) strongly depend on the spine morphology, which is highly variable for pyramidal cells. Different currents and diameters of idealized neck and head configurations are compared. Spine A corresponds to the spine shown in Fig. 1. In smaller spine heads the concentration changes are stronger than in larger spine heads. A thinner spine neck also increases the depolarization and concentration changes. The total electric resistance of the spine *R*_*e*_ increases as a result of the concentration changes.

### Diffusion constants are relevant to accurately estimate the electrical resistance

As a result of the concentration changes, a previous modelling study based on PNP-equations predicted that the intracellular resistance decreases during synaptic input [17]. Therefor, we compute the total resistance 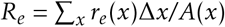 by taking the sum over all finite segments along the spine (compare Fig. 4G). *r*_*e*_ denotes the resistivity and *A* the cross section (see Methods section). However, when we analyze the total resistance along the spine during current injection, we find the resistance slightly increasing instead (Fig. 2, last row). A major difference between both studies is that we consider sodium, potassium and chloride as different ion-species at physiological ion concentrations and with different diffusion constants while [17] distinguish only between positive and negative ions and assume that both ion types have the same concentration and diffusivity.

To test whether we could replicate the findings in [17], we repeated the simulations of Figure 2, but with altered parameters. As in [17], we set the diffusion constant of sodium to be equal that of potassium and chloride. Interestingly the cumulative resistance is now decreasing during current injection (Fig. 3). Next, we increased the concentration of chloride to 150 mmol. Positive and negative ions now have the same concentration and diffusivity. Here, we found that the decrease of the resistance became even more prominent (Fig. 6) and comparable to [17].

**Figure 3:**
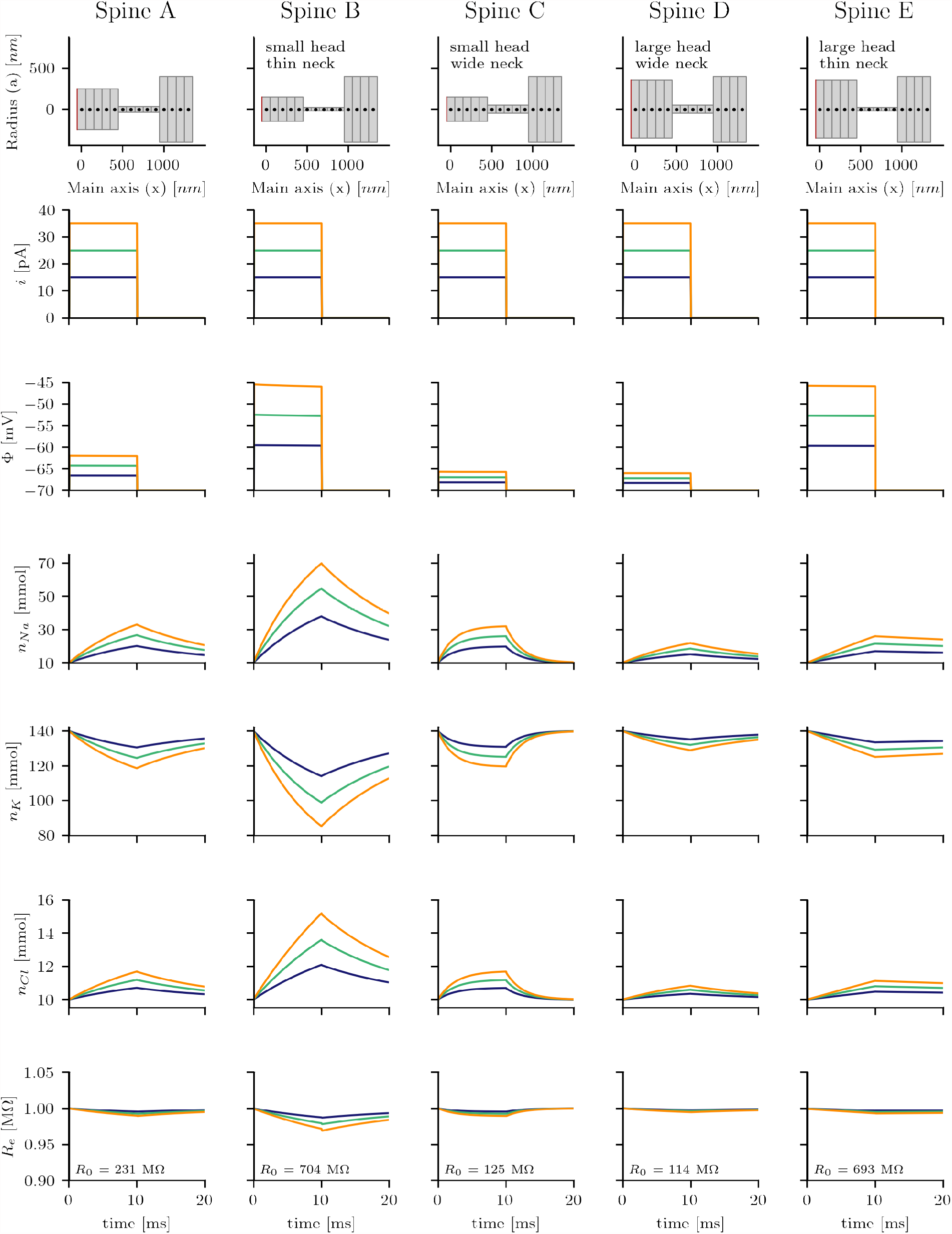
The simulations of Fig. 2 are repeated with a higher diffusivity of sodium. The diffusion constants of sodium, potassium and chloride are now identical. The membrane depolarization is set equal to the product of the total resistance *R*_*e*_, and the strength of the injected current *i*. But the total resistance *R*_*e*_ now decreases instead.

**Figure 4:**
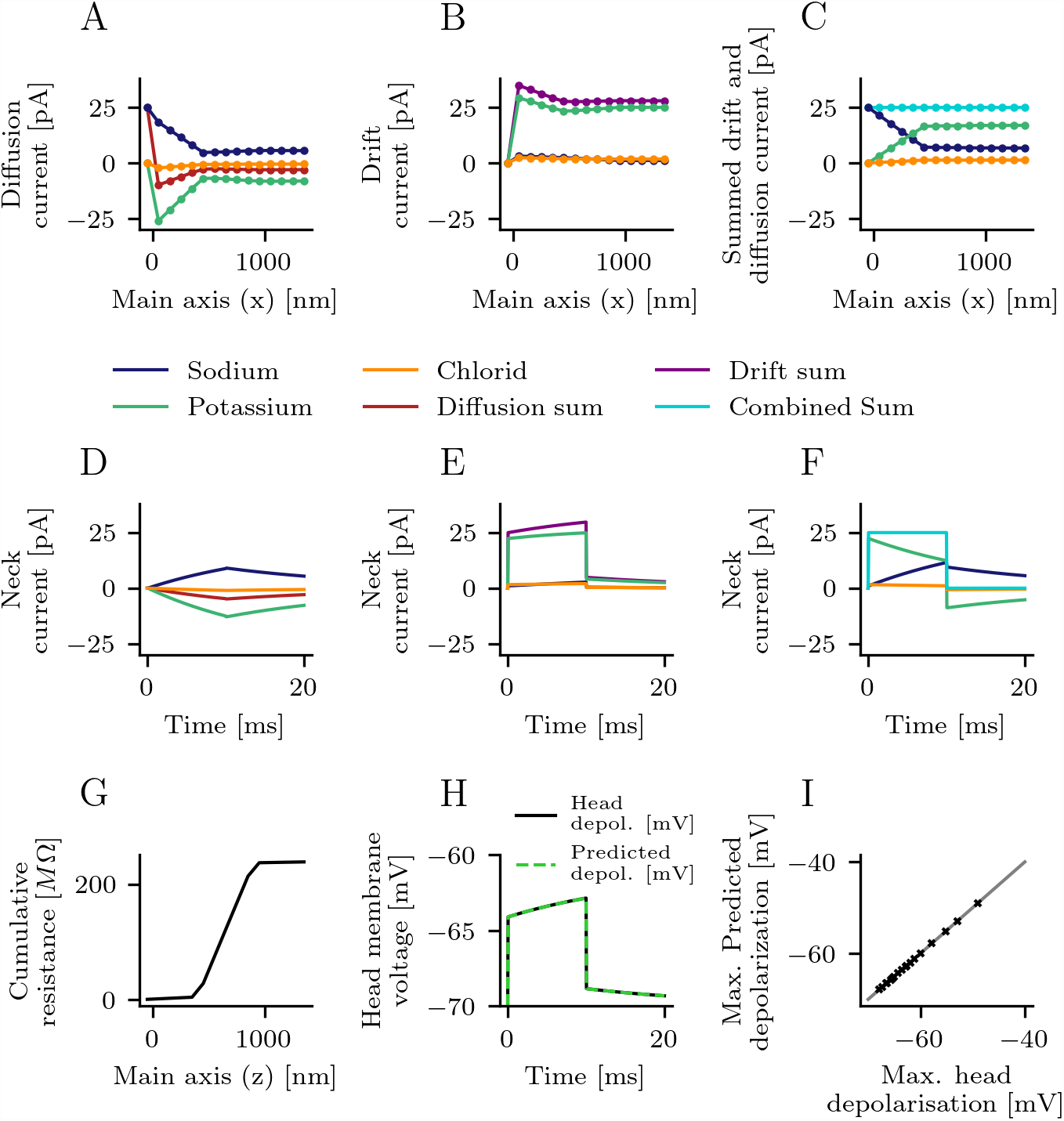
The electric currents of the simulation shown in Fig. 1 are broken down. A) Electric currents evoked by the diffusion of ions between the lattice points after 10 ms of current injection (25 pA). B) Electric currents evoked by the electric field (drift current) after 10 ms of current injection. C) Sum of the currents shown in A) and B). The total electric current (sum of drift and diffusion currents) exactly matches the injected current everywhere in the spine. An increased electric field therefore compensates the diffusion of electric charges. D) Diffusion currents through spine neck. E) Drift currents through spine neck. F) Summed current through spine neck exactely matches the injected current. G) The cumulative resistance along the spatial lattice after 10 ms. Almost all of the resistance is caused by the spine neck. H) The product of the total current evoked by the electric field (purple line in B) and the total resistance (max. value of cumulative resistance as shown in G) theoretically predict the measured membrane voltage in the spine head. Diffusion currents, therefore increase the membrane voltage in spines. I) Identical to the case shown in H) the theoretically predicted depolarization and the measured depolarization after 10 ms match for all simulations shown in Fig. 2.

This effect can be explained as follows: In our original setup sodium has a 35 % reduced diffusivity compared to potassium. During current injection the sodium concentration increases while the concentration of potassium drops. As the electrical resistance is inversely proportional to the diffusion constants of the ions it has to decrease. In the second setup, where all diffusion constants are equal, the total number of ions in the spine is increasing even though the membrane potential does not change. To compensate injected sodium, cations, predominantly potassium cations, leave the spine. But chloride anions also enter the spine. The total charge remains unchanged but the number of ions is increasing. This effect gets stronger when more chloride ions are available in the system.

We conclude that the right choice of diffusion constants and ion concentrations is essential to reliably study the electric function of denritic spines.

### Diffusive currents boost spine depolarisation

As shown before the spine head’s membrane potential is continuously increasing during current injection (Fig. 2, second row). This effect cannot be explained by the small changes of the intracellular resistance (Fig. 2, last row). To understand how the head’s membrane potential is changing, we analyze the electrical currents in detail. These currents can be separated into currents caused by the electric field and currents caused by ion-concentration gradients.

As shown in Figure 1E-G and Figure 2 the ion concentrations are considerably changing in the spine head during current injection. Along with the increasing concentration gradients, this increases the diffusion of ions and leads to an measurable electrical current (diffusion current). Indeed, after 10 ms of current injection we find considerable diffusion currents (Fig 4A). Currents in the head are larger because of the larger radius a. Sodium and chloride ions are diffusing out and potassium ions are diffusing into the spine head. In total, the sum of all diffusion currents is negative. Next (Fig. 4B), we looked at currents along the spine’s main axis induced by the electric field (drift current). All drift currents are positive, so that, on average, the electric field forces sodium potassium ions to leave the spine head and chloride ions to enter the spine head. Most of the drift current is caused by potassium ions which have the highest concentration of all ions in the intracellular space. Interestingly, the summed electric current is higher than the injected current of 25 pA. But when we compute the sum of all currents (Fig. 4C) we find that diffusion and drift currents exactly add up to 25 *pA* everywhere in the spine, so that they are equal to the injected current. In summary, as the ion concentrations are changing, this leads to diffusional currents. To maintain a balance between positive and negative charges the electrical currents have to compensate the diffusional currents.

Next we study the different currents through the spine neck over time. As the concentration gradients between head and dendrite are increasing the diffusion currents get stronger, but have a negative value (Fig. 4D). The diffusion currents persist after the end of the current injections at 10 ms. The drift currents are increasing over time. A small drift current continues after the end of the current input. The total neck current always matches the injected current exactely (Fig 4F). Next, we analyze the cumulative electrical resistance of the simulated spine (Fig. 4G). Most of the total resistance is caused by the spine neck, but the resistance is only slighly changing over time (compare Fig 2) and can not explain the changes of the head depolarization. However, when we estimate the voltage drop across the simulated spine from the electrical resistance and the drift current *I*_*e*_ as 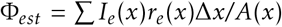, instead of the total current (Fig 5B,E) we can accurately predict the spine head voltage (Fig. 5H). To verify this for other spines, we compare the maximum value of the predicted head voltage with the maximum value of the measured head membrane potential (Fig 4I). Both values agree for all spines and input currents shown in Figure 2.

**Figure 5:**
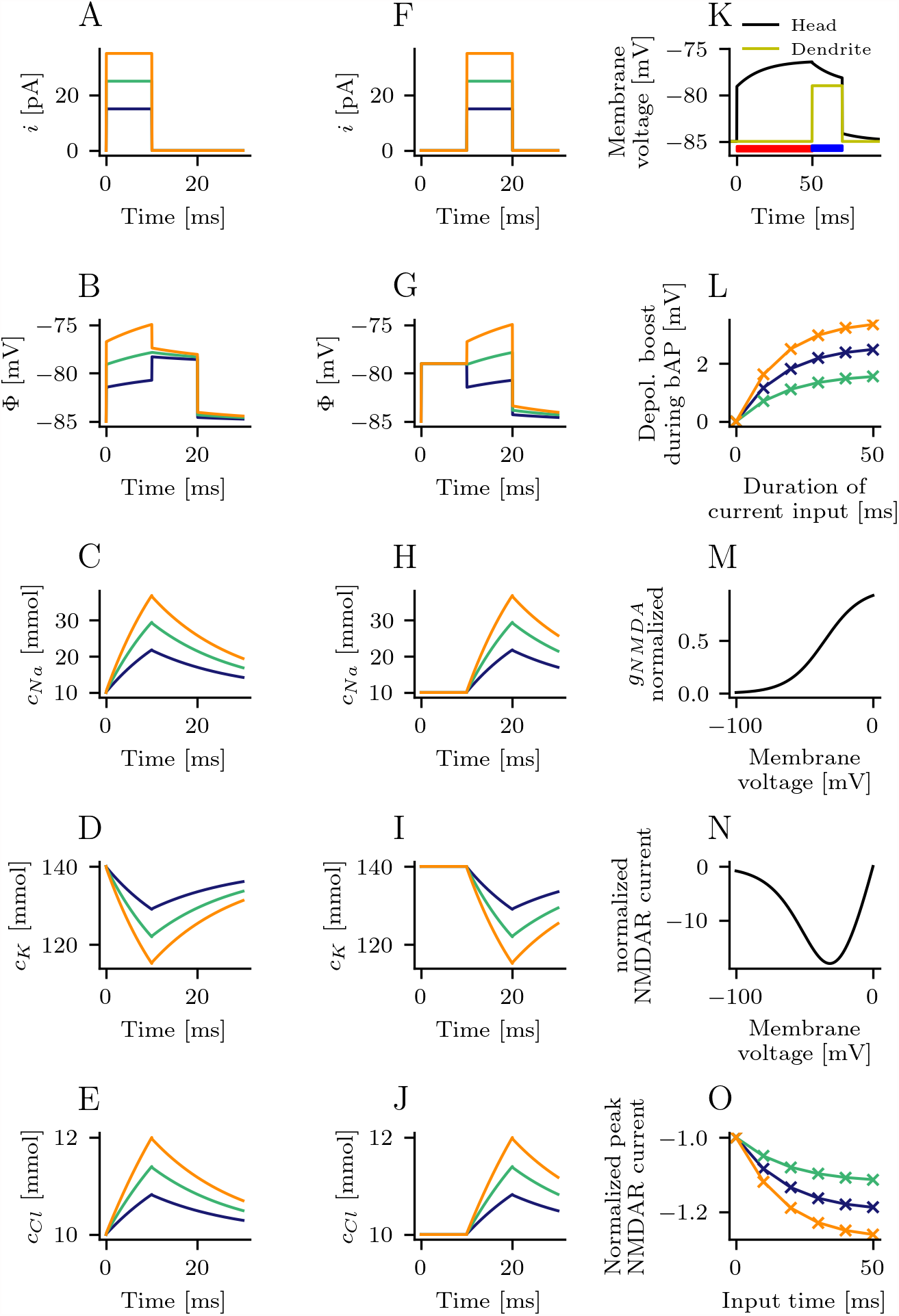
A-E) The electric potential and the ion concentrations in the spine head when a current is injected for 10 ms and then the dendrite gets depolarized by 6 mV for the next 10 ms. F-J) The electric potential and the ion concentrations in the spine head when the dendrite gets depolarized by 6 mV for 10 ms and then a current is injected for the next 10 ms. K) A variable duration (red bar) of the injected current is used to mimic variable presynaptic activity. Membrane potential and calcium influx are analyzed in the subsequent phase of postsynaptic activity (blue bar). L) Increasing the duration and the strength of the current injection boosts the spine head’s deploarization. The shown voltage will add to the 6 mV of dendritic depolarization. M) Conductivity of NMDARs grows as a function of voltage. N) Normalized NMDAR-current as function of holding potential. O) Peak NMDAR-current during the phase of dendritic depolarization due to a back-propagating action potential. The increased membrane voltage amplifies the calcium influx. M)-O) The model of the channel conductance *g*_*NMDA*_ and the NMDAR current are reproduced from [8].

**Figure 6:**
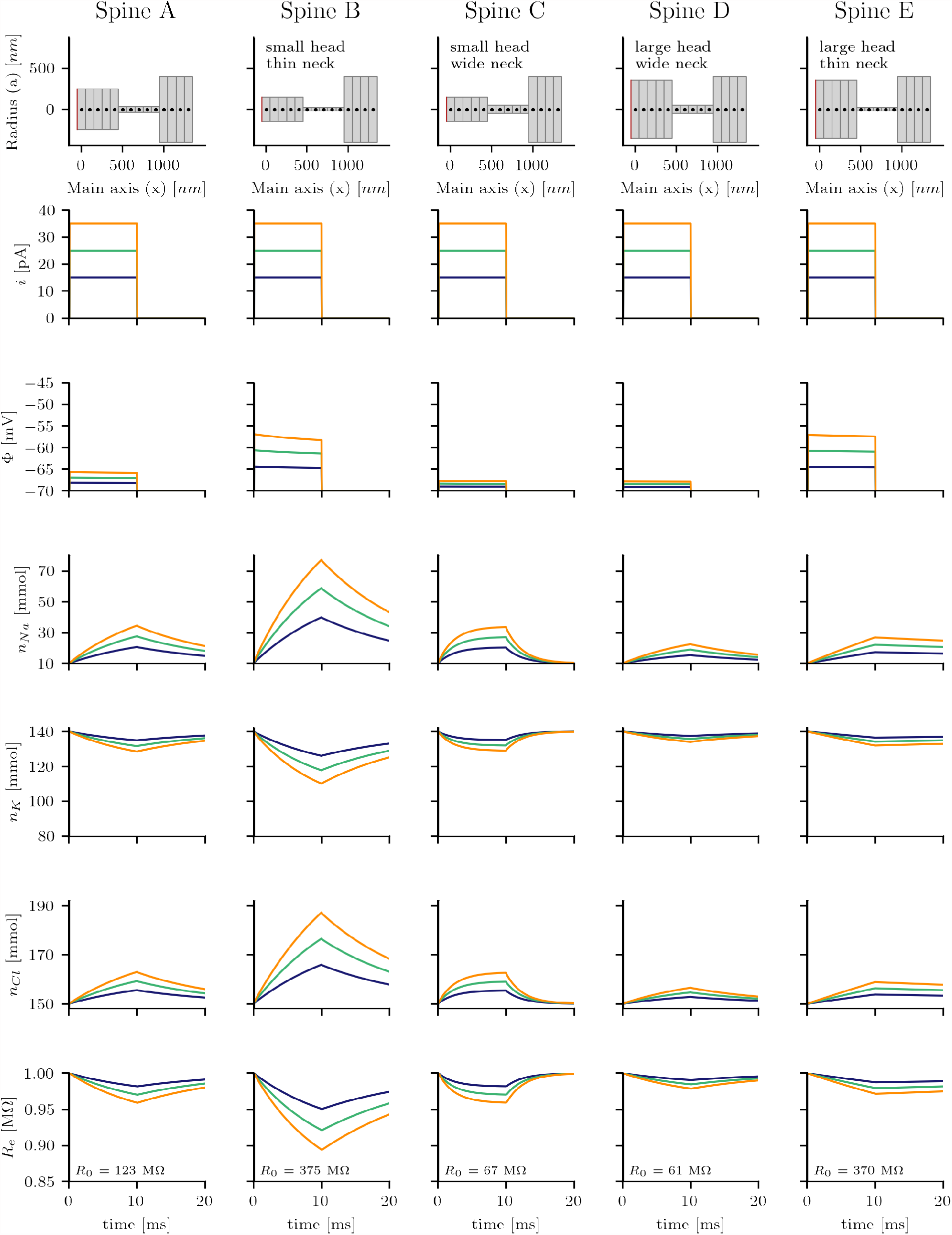
The simulations of Figure 02 and 03 are repeated again. The diffusion constants of sodium, potassium and chloride are identical. The resting concentration of chloride is set to 150 mmol. The number of free anions and free cations is now identical. The discrepancies between the simulation results of Fig. 02 and 03 get even stronger.

In summary, ion-concentration gradients lead to diffusion currents. These diffusion currents are compensated by a change in the electrical drift currents which in turn increase the membrane potential.

### Interaction between synaptic current injection and dendritic depolarization

The coincidence of pre and postsynaptic action potential can trigger synaptic plasticity in dendritic spines [29]. The direction of the changes in synpatic strenght are determined by the temporal order of pre- and postsynaptic activity. The magnitude is positively correlated with the peak elevation of the intracellular *Ca*^2+^ concentration [22]. The main pathway for *Ca*^2+^ influx from the extracellular space into dendritic spines is through NMDA-type glutamate receptors (NMDARs). The channel conductance of NMDARs is, via the *Mg*^2+^ block, highly voltage dependent [4]. But the membrane voltage in spine heads gets boosted, as we found before, by ion diffusion through the spine neck after current injection. Therefore we ask how the strength, duration and the timing of synaptic input could affect the NMDAR-channel mediated current during sequences of pre- and postsynaptic action potentials. In the following section the parameters of NMDAR kinetics and the resting potential are based on [8]. The other spine parameters are identical to those of the simulation shown in Fig. 1.

To test whether the spine depolarization depends on the chronological order of pre- and post-synaptic action potential, we compared two cases. In the first case a current (15 - 35 pA) is injected for 10 ms into the spine head (Fig. 5A). Subsequently the dendritic end gets depolarized by 6 mV for 10 ms (Fig 5B). In the second case the dendritic end is depolarized first (Fig. 5G), followed by a current injection into the spine head (Fig.5F). In the first case the ion concentrations are alterd after 10 ms of current injection (Fig. 5C,D,E). The diffusion currents persist into the phase of dendritic depolarization and raise the membrane voltage of the spine head above the dendritic depolarization of 6 mV (Fig. 5B). Differently in the second case; the depolarization of the dendrite propagates into the spine head without attenuation (Fig. 5G). The ion concentrations are unaffected and remain constant inside the spine head during the phase of dendritic depolarization (Fig. 5H,I,J). There are no lasting effects for the following time of current injection.

Therefore, we focus on sequences where a synaptic input is followed by a postsynaptic action potential. The boost of the spine head’s membrane voltage during postsynaptic activity depends on the synaptic input. To quantify this dependency a current of variable strength and duration is injected (red bar in Fig 5K indicates duration of current injection). Right after the current injection, the dendrite gets depolarized by 6 mV for 10 ms again (blue bar in Fig. 5K). It can be seen that when the duration of current injection is increased from 10 ms to 50 ms the depolarization boost increases from 0.70 mV to 1.55 mV for 15 pA current strength and from 1.62 mV up 3.34 to mV for 35 pA current (Fig. 5L). Based on 6 mV depolarization in the dendrite the heads membrane voltage gets amplified by up to 55.6 % in the tested cases.

The NMDAR channel conductance increases as a function of the membrane voltage (Fig 5M). This leads to a higher NMDAR current in the relevant votage range (Fig. 5N). Therefor we compute the peak NMDAR current as function of the input strength and duration. We find that current through NMDARs can be boosted by roughly up to 25% for realistic spine parameters. In summary, concentration changes caused by synaptic input amplify the spine head voltage and the calcium influx during subsequent postsynaptic activity.

## 4 Discussion

Dendritic spines have a very small volume. As a consequence, physiological electric currents can alter the ionic composition of the spine’s intracellular space. The resulting concentration gradients lead to additional electric currents caused by the diffusion of ions. These currents are not captured by the cable equation [25], as it only considers the electric field as driving force of ion movement. The PNP-equations on the other hand are capable of accurately estimating the ion-movement in spines based on electrodiffusion. But due to the high complexity of this equation system, they are not suited to find solutions in larger systems with multiple ion species. The system of equations used here closes the gap between the two formalisms. It considers the electric potential and time-dependent ion concentrations of an arbitrary number of different ions. Still, basic numerical algorithms can be used to find solutions. In this study we solve the equations using finite differences and a simple solver with explicit time stepping. The derivation of similar equations was done before [14], but to our knowledge they have not been used to study electrodiffusion within spines.

In this study we simulate ionic currents in dendritic spines during current injection and find that the ion concentrations in dendritic spines are considerably changing. When spine parameters are chosen in agreement with experimental findings [10], the sodium concentration increases by almost 300% after 10 *ms* of a 25 *pA* input current. This is also in agreement with previous simulation studies based on PNP-equations [17, 24]. The concentration gradients and the resulting diffusion of ions lead to additional currents in the spine. The summed current evoked by diffusion flows in the opposite direction compared to the current evoked by the electric field. To maintain the balance between positive and negative charges in the spine head, the electric field increases and compensates the additional diffusion of ions. As a result the depolarization of the spine head rises by approximately 20 % in this case.

To accurately estimate all ionic currents, the correct choice of the diffusion constants and ionic concentrations is critical. Sodium has a lower diffusivity than chloride and potassium. And the number of freely moving anions (chloride) is lower than the number of cations (sodium and potassium). If that is not taken into account the results can be complementary. In systems with equal diffusion constants, the concentration gradients will not boost the head’s membrane potential and predict an opposite change in the intracellular resistance [17]. Changes in the chloride resting concentration increase these discrepancies. To better understand this phenomenon, one can relate it to the Goldman-equation. Here, the membrane potential evoked by sodium, potassium and chloride depends on the exact ionic concentrations on both sides of the membrane and the different permeabilities of the three ion species inside the membrane. The permeability can be compared to the neck resistance for each ion, and the intracellular and extracellular ion concentrations correspond to head and dendrite ion concentrations. An important difference, however, is that the effective current across the membrane is zero in the Goldman equation, while here it is equal to the strength of the current injection.

It was found that changes in the synaptic plasticity can be triggered by a pairing of pre- and postsynaptic action potentials. Thereby the temporal order is important to set the direction of changes in the synaptic strength. Postsynaptic activity that follows synaptic input within a short time can lead to long-term potentiation of the synapse. The peak elevation of C a2+ concentration controls the magnitude of the changes [22]. Here we identified a previously unknown mechanism that will increase peak calcium levels in spines in cases when postsynaptic activity follows a synaptic input. The peak concentration depends on the strength of the current input. As shown by our simulations, a current injection leads to changes in the ionic concentrations that are still present during a subsequent depolarization of the dendrite. The currents conditioned by the diffusion of ions amplify the dendritic depolarization in the spine head. This amplified membrane depolarization in the head increases the conductance of NMDAR-channels and thereby elevates the calcium influx into the spine. In average pyramidal cell spines, we found that this effect can increase peak calcium influx by up to 25 % for strong current inputs, as expected during repeated activation of the synapse, e.g., during a presynaptic spike train. This effect depends on the spine’s morphology and is especially important for small spines with long and thin necks. Our observation indicates that different mechanisms for learning and electrical function could be at work in these spines.

The equations presented here can be considered as a generalization of the cable equation. By setting the ion concentrations to a constant value, one recovers the well-known form of the cable equation. By discretizing the cable along the main axis into compartments of finite size one can build multicompartment models to study large dendritic trees [13]. The same concept can be applied to the equations presented here. This can be utilized to extend compartmental models and to accurately estimate synaptic currents including the diffusion of ions. Finally, the inclusion of larger molecules into the model, such as dye molecules used for photo stimulation, might also help to correctly interpret experimental results from imaging spines.

## Acknowledgements

I am deeply grateful to my professor and supervisor, Prof. Dr. Andreas V. M. Herz, for his guidance and support throughout my doctoral studies. Furthermore, I would like to acknowledge the generous support of the Ludwig-Maximilians Universität, the German Federal Ministry of Education and Research, and the Bernstein Center for Computational Neuroscience Munich, who provided the funding necessary to carry out this research.

